# The development of compulsive coping behaviors depends on the engagement of dorsolateral striatum dopamine-dependent mechanisms

**DOI:** 10.1101/2022.09.27.509680

**Authors:** Chiara Giuliano, Lucia Marti-Prats, Ana Domi, Mickaël Puaud, Yolanda Pena-Oliver, Colin McKenzie, Barry J. Everitt, David Belin

## Abstract

Humans greatly differ in how they cope with stress, a natural behavior learnt through negative reinforcement. Some individuals engage in displacement activities, others in exercise or comfort eating, and others still in alcohol use. Across species, adjunctive behaviors, such as polydipsic drinking, are used as a form of displacement activity that reduces distress. Some individuals, in particular those that use alcohol to self-medicate, tend to lose control over such coping behaviors, which become excessive and compulsive. However, the psychological and neural mechanisms underlying this individual vulnerability have not been elucidated. Here we tested the hypothesis that the development of compulsive adjunctive behaviors stems from the functional engagement of the dorsolateral striatum (DLS) dopamine-dependent habit system after a prolonged history of adjunctive responding. We measured in longitudinal studies in male Sprague Dawley rats the sensitivity of early established vs compulsive polydipsic water or alcohol drinking to a bilateral infusion of the dopamine receptor antagonist α-flupentixol into the anterior DLS (aDLS). While most rats acquired a polydipsic drinking response with water, others only did so with alcohol. Whether reliant on water or alcohol, the acquisition of this coping response was insensitive to aDLS dopamine receptor blockade. In contrast, after prolonged experience, adjunctive drinking became dependent on the aDLS dopamine-dependent habit system at a time it was compulsive in vulnerable individuals. These data suggest that habits may develop out of negative reinforcement and that the engagement of their underlying striatal system is necessary for the manifestation of adjunctive behaviors.

**Significance statement:** Harnessing the individual variability that rodents, like humans, show to engage in adaptive or maladaptive coping strategies, which can result in the development of compulsive disorders, here we demonstrate that the functional engagement of the dorsolateral striatum-dependent habit system precipitates the transition to compulsion in rats that have acquired a polydipsic adjunctive drinking response with water or alcohol as a means to cope with distress. The results of this study not only provide evidence for the emergence of instrumental habits under negative reinforcement, but they also reveal that compulsive behaviors that originate from the loss of control over coping strategies are mediated by the dorsolateral striatum-dependent habit system.

## Introduction

When facing the distress generated by a challenging, emotionally taxing, aversive situation, individuals greatly differ in the emotion regulation strategy they use, which is influenced by situational demands (1-6). Emotion regulation and associated coping strategies (7), such as exercise, comfort eating, shopping, or displacement behaviors (8, 9), which all decrease stress through negative reinforcement, have long been suggested to be a prerequisite for adaptive functioning, the promotion of resilience and well-being (10, 11).

Several species cope with stress using a form of displacement called adjunctive behaviors (12, 13). One such adjunctive anxiolytic response, schedule-induced polydipsia (SIP) (12-15), manifests itself as polydipsic water intake in the face of intermittent food delivery in food-restricted animals (16-18). At the population level, non-regulatory polydipsic drinking develops over a week and remains stable for long periods of time during which it selectively decreases the levels of stress-related hormones that had been increased by the associated intermittent food delivery (14, 15, 19-22).

However, some humans (23-30) and individuals of other species (31) characterized, for instance by a high impulsivity trait (32), lose control over these coping strategies which become excessive and promote the development of compulsive disorders such as obsessive compulsive and substance use disorders.

As exemplified by the emotion regulation challenges the COVID-19 pandemic has posed (33-36), under circumstances perceived as overwhelming, some individuals resort to alternative means to cope with stress, such as the use of alcohol (37-42) or other drugs (43), a strategy associated with a greater vulnerability to develop compulsivity (42, 44-52). Across species the individual tendency to rely on the anxiolytic properties of alcohol to cope with distress (44, 52, 53), including that induced by the SIP procedure (54), has been associated with an increased vulnerability to switch from controlled to compulsive alcohol use, a key feature of alcohol use disorder (AUD) (44, 51, 52, 54-56).

However, the psychological and neural basis of the individual vulnerability to lose control over coping behaviors, involving alcohol use or not, and the ensuing development of compulsivity have not been defined. Increasing evidence suggests that the transition from controlled goal-directed behaviors to compulsion, including the compulsive seeking and drinking of alcohol (57, 58) and the development of compulsive adjunctive polydipsic drinking, is dependent on a shift in the locus of control over behavior from the ventral to the dorsolateral striatum (DLS)-dependent habit system (59). Thus, while the reinforcing properties of alcohol mediated by the mesolimbic system support recreational alcohol use (60, 61), it is the engagement of anterior DLS (aDLS) dopamine (DA) dependent alcohol seeking habits that promotes the transition to compulsive alcohol seeking and drinking (57, 58, 61). Similarly, the development of adjunctive polydipsic water drinking behavior, but not its compulsive manifestation, is dependent on the mesolimbic system (62, 63) while well-established excessive, compulsive adjunctive water drinking in vulnerable individuals has been associated with an increased in spine density in DLS medium spiny neurons (64).

Here we tested the hypothesis that the development of compulsive coping behavior, as manifested as excessive polydipsic drinking, whether of alcohol or water, is associated with functional engagement of, and an increased reliance on, the DLS-DA dependent habit system. To test this hypothesis, we assessed the sensitivity of the polydipsic water or alcohol drinking response of each individual in large cohorts of outbred rats to bilateral aDLS DA receptor blockade (54, 58, 65) over the course of the acquisition of each coping behavior and the subsequent transition to compulsivity.

## Methods and Materials

### Subjects

Ninety-four male Sprague Dawley rats (Charles River, UK), from three different cohorts, weighing 300-350g at the start of the experiments, were used in this study. Rats were single-housed under a 12-hour reversed light/dark cycle (lights off at 7:00 AM) and food restricted to gradually reach 85% of their theoretical free-feeding body weight before the start of the behavioral training. Water was always available *ad libitum*. Experiments were performed 6-7 days/week between 8 am-5 pm. All experimental protocols were conducted under the project license 70/8072 held by David Belin in accordance with the regulatory requirement of the UK Animals (Scientific Procedures) Act 1986, amendment regulations 2012, following ethical review by the University of Cambridge Animal Welfare and Ethical Review Body (AWERB).

### Experimental procedures

The series of experiments conducted in this study are schematically summarised in **Figure 1**.

**Figure 1:**
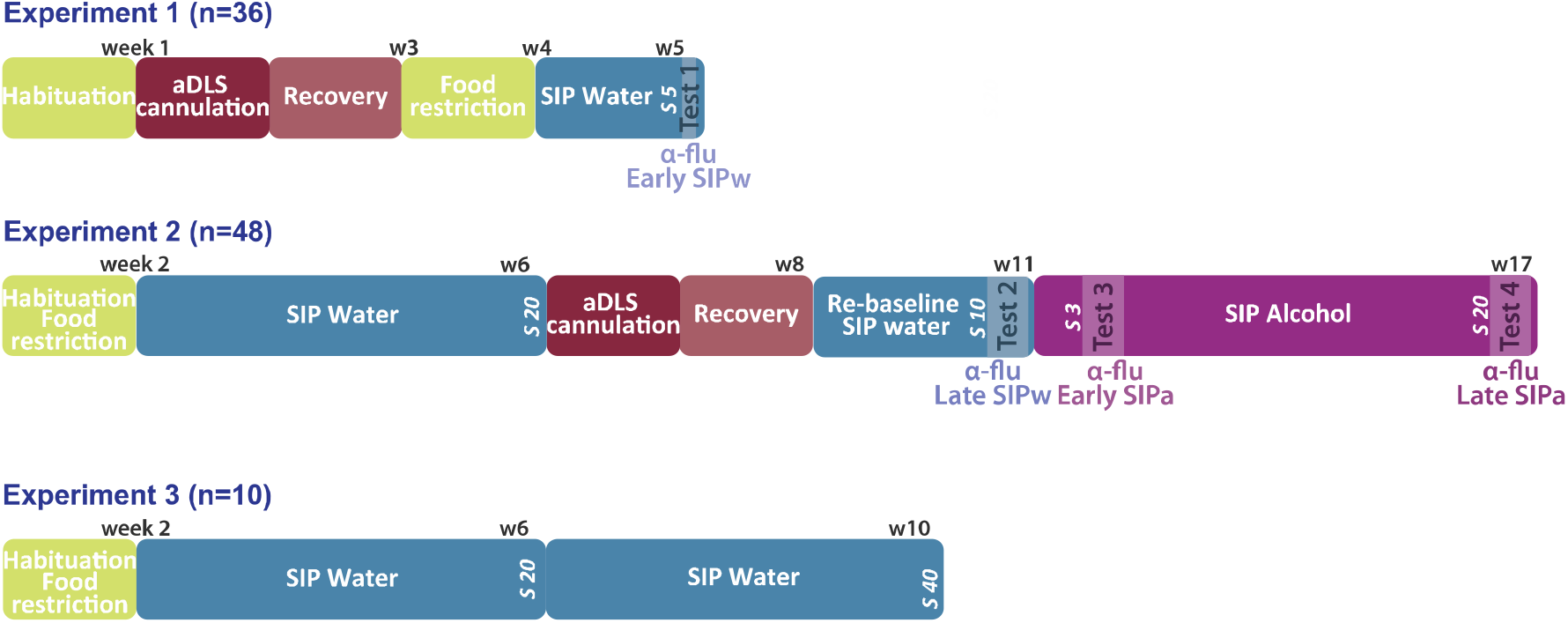
Timeline of the experiments. Timeline of the three experiments carried-out on independent cohorts of male Sprague Dawley rats. **Experiment 1:** Following a week of habituation to the animal facility, thirty-six rats received bilateral cannulation of their anterior dorsolateral striatum (aDLS). A week later, rats were food restricted to 85% of their theoretical free-feeding weight and trained in a schedule-induced polydipsia (SIP) procedure with water (SIPw). The reliance of the acquisition of adjunctive water drinking on aDLS dopaminergic mechanisms was assessed after 5 SIPw sessions as the sensitivity of drinking behavior to bilateral infusion of the dopamine (DA) receptor antagonist α-flupentixol (α-flu, 0, 6, 12 μg/side, between-subject design) into the aDLS (**Test 1**, α**-flu Early SIPw**). **Experiment 2**: Following a week of habituation to the animal facility, forty-eight rats were food restricted for a week before being trained in the SIP procedure with water for 20 sessions, e.g., until the establishment of aberrant levels of water intake in vulnerable individuals. Then, rats were implanted with bilateral cannulas targeting the aDLS and, after at least a week, they were re-baselined under SIPw for 10 sessions. Then, the reliance of well-established, compulsive, adjunctive water drinking behavior on aDLS DA was assessed as the sensitivity of water drinking behavior to bilateral infusion of α-flupentixol (0, 6, 12 μg/side, within-subject design) into the aDLS (**Test 2**, α**-flu Late SIPw**). Subsequently, water was replaced by 10% alcohol (SIPa) and the reliance of early and well-established adjunctive alcohol drinking on aDLS DA was measured after 3 (**Test 3**, α**-flu Early SIPa)** or 20 (**Test 4**, α**-flu Late SIPa**) SIPa sessions, respectively. **Experiment 3**: In order to test the stability of water intake levels once adjunctive water drinking has been established, ten rats were food restricted after a week of habituation to the colony, and then trained in the SIP procedure for 40 sessions, a period of training similar to that used in experiment 2, but with water as the only available solution throughout. W, week; s, session.

The first experiment aimed to determine the involvement of aDLS DA-dependent mechanisms (referred to subsequently as aDLS DA) in the acquisition of a coping adjunctive water drinking response. Thus, after a week of habituation to the vivarium, thirty-six rats received bilateral cannulations of the aDLS and, following recovery, were food restricted to progressively reach 85% of their theoretical free-feeding weight. Rats were then trained in the SIP procedure with water (SIPw). Following one habituation and one baseline session (see SIP section below), rats were exposed to five 60-min SIPw sessions before the sensitivity of their early adjunctive water drinking to aDLS DA receptor blockade was measured (**Test 1, Effect of** α**-flu on Early SIPw**). Fourteen rats were excluded from the experiment because of loss of their guide cannula or cannula misplacements, so that twenty-two rats were included in the final analysis.

The second experiment aimed to test the reliance on aDLS DA of well-established adjunctive water drinking vs that of early and well-established adjunctive alcohol drinking. Thus, following a week of habituation to the vivarium, forty-eight rats were food restricted to progressively reach 85% of their theoretical free-feeding weight before starting behavioral training. Following one habituation and one baseline session (see SIP section below), rats were exposed to twenty 60-min SIPw sessions, e.g., until the establishment of aberrant, compulsive, levels of water intake in vulnerable individuals (31, 32, 54, 66). Rats were subsequently implanted bilaterally with intra aDLS cannulae and, following recovery, they were re-baselined under SIPw for ten sessions before the reliance of their well-established adjunctive water drinking on aDLS DA was measured (**Test 2, Effect of** α**-flu on Late SIPw**). Then, water was replaced by 10% alcohol and rats were trained to maintain their adjunctive behavior now using alcohol (SIP with alcohol, SIPa), or to acquire the coping response for those that rely on alcohol to cope with distress, as previously described (54). The reliance of early and well-established adjunctive alcohol drinking on aDLS DA was measured after three (**Test 3, Effect of** α**-flu on Early SIPa**) or twenty daily sessions (**Test 4, Effect of** α**-flu on Late SIPa**), respectively. Because ten individuals lost their cannulae or had misplaced cannulae as revealed after post-mortem histological assessment, thirty-eight rats were included in the final analysis.

The third experiment aimed to test the stability of polydipsic water intake levels over a period of training in the SIP procedure similar to that of rats in experiment 2. Thus, after a week of habituation to the vivarium, ten rats were food restricted to progressively reach 85% of their theoretical free-feeding weight before starting behavioral training. Following one habituation and one baseline session (see SIP section below), rats were exposed to forty 60-min SIPw sessions.

### Schedule-induced polydipsia (SIP)

#### Apparatus

SIP training was carried out as previously described (32, 66, 67) in twelve operant chambers located within ventilated sound-attenuating cubicles (Med associates, St. Albans, USA) and made of aluminium and transparent acrylic plastic with a stainless-steel grid floor (24 cm x 25.4 cm x 26.7 cm). Chambers were equipped with a house light (3-W), a food tray (magazine) installed at the centre of the front wall, and a bottle from which a stainless-steel sipper tube delivered water or alcohol into a receptacle placed in a magazine on the wall opposite the food magazine. Water or alcohol were freely available throughout the 60-min sessions.

#### Behavioral training

SIP training consisted of the following stages:

#### Habituation / baseline water intake

During the habituation session, rats were given access to water and sixty food pellets (45 mg, TestDiet, USA) that had been placed in the magazine. The volume of water consumed by each rat over this 60-min session was measured in order to determine the amount of water each individual drank to meet their homeostatic needs while eating sixty 45 mg pellets. Rats were then tested in one 60-min magazine training session during which sixty food pellets were delivered under a variable interval 60-second schedule (VI-60 s) over 60 min. This magazine training provided the baseline level of homeostatic water intake over one hour.

#### SIP with water (SIPw)

The SIP procedure was based on a fixed-time 60-second (FT-60 s) schedule of food delivery, previously shown to induce adjunctive drinking behavior with robust and persistent individual differences in the tendency to develop excessive, compulsive adjunctive drinking behavior (32, 54, 66, 68, 69).

Twenty-four hours after the baseline session, rats underwent five (Experiment 1), twenty (Experiment 2) or forty (Experiment 3) of these 60 min FT-60 s SIPw training sessions. 300 mL bottles filled with fresh water were weighed and inserted into the operant box immediately prior to the initiation of each session. House lights were switched on at the beginning and switched off at the end of each session. The total amount of water consumed (mL) was calculated daily as the difference between the weights of the bottle before and after the session.

Individuals whose average water consumption over the last three days of training was in the upper and lower quartiles of the population were considered as high (HD) and low drinkers (LD), respectively, as previously described (32, 54, 66).

#### SIP alcohol (SIPa)

Twenty-four hours after the last SIPw session, water was replaced by 10% alcohol and rats were trained for twenty 60-min SIPa sessions. The total amount of alcohol consumed (mL) was calculated daily as the difference between the weights of the bottle before and after the session.

### Drugs

The alcohol solution was prepared by mixing 99.8% ethanol (Sigma-Aldrich, UK) with tap water to obtain 10% alcohol (54, 67).

The DA receptor antagonist α-flupentixol (Sigma-Aldrich, UK) was dissolved in double-distilled water as previously described (70). Drug doses (6 or 12 μg/side), reported in the salt form were selected to be in the range of those previously shown effectively to decrease aDLS-dependent cocaine, heroin or alcohol seeking behavior (58, 70-72).

### Surgery: aDLS cannulations

Rats underwent stereotaxic surgery either before behavioral training (Experiment 1), or after acquisition of SIPw (Experiment 2) under isoflurane anaesthesia (O2: 2L/min, 5% isoflurane for induction and 2 % for maintenance), as previously described (58). Guide cannulae (22-gauge, Plastics One, Roanoke, VA, USA) were bilaterally implanted 2 mm above the aDLS (anterior/posterior (AP) +1.2, mediolateral (ML) ±3, dorsal/ventral (DV) -3; AP and ML coordinates measured from bregma, DV coordinates from the skull, incisor bar at −3.3 mm), as previously described (58, 73). Cannulae were held in place using dental acrylic cement anchored to four stainless steel screws tapped into the frontal and parietal bones of the skull. Obturators (Plastics One, Roanoke, VA, USA) were placed in the cannulae to maintain patency. All animals were given five days to recover from surgery and for the first three days after surgery, rats were treated daily with an analgesic agent (1 mg/kg Metacam, Boehringer Ingelheim, Ingelheim am Rhein, Germany) orally administered in drinking water.

### Intra-striatal infusions

The influence of aDLS DA receptor blockade on adjunctive drinking behavior was tested at early and late stage of training for SIPw and SIPa, namely during the acquisition of SIPw (Experiment 1, SIPw session 5 onwards, **Test 1 Effect of** α**-flu on Early SIPw**) and SIPa (Experiment 2, SIPa session 3 onwards, **Test 3, Effect of** α**-flu on Early SIPa**) and well-established SIPw (Experiment 2, SIPw session 20 onwards, **Test 2, Effect of** α**-flu on Late SIPw**) and SIPa (SIPa session 20 onwards, **Test 4, Effect of** α**-flu on Late SIPa**).

Before being tested, rats were habituated to the intra-aDLS insertion of the injector. Each test was preceded by intra-aDLS infusions (0.5 μl/side) of α-flupentixol (0, 6, 12 μg/side, made via 28-gauge steel injectors (Plastics One, Roanoke, VA, USA) lowered to the injection sites 2 mm ventral to the end of the guide cannulae. Infusions were made over 90 s using a syringe pump (Harvard Apparatus) and were followed by a 60 s period to allow diffusion of the infused drug or vehicle before injectors were removed and obturators were replaced. Test sessions began 5 min later. The effect of aDLS DA receptor blockade on adjunctive drinking was tested on rats from experiment 1 (**Test 1**) in a between subject design, and in rats from experiment 2 (**Tests 2, 3 and 4**) in a counterbalanced order following a Latin-Square design. Each infusion-day was followed by two baseline sessions.

### Histology

At the end of the experiment, rats were euthanized with an overdose of sodium pentobarbital (300 mg; Dolethal; Vétoquinol UK Ltd, Buckingham, UK) and then perfused transcardially with isotonic saline followed by 10% neutral buffered formalin. Brains were extracted and transferred to a 30% sucrose solution in 0.01 M PBS for 48 hours before being processed into 60 μm coronal sections using a Leica CM3050 S Research Cryostat. Sections were mounted and stained with Cresyl Violet.

Cannulae placements in the aDLS were verified using a light microscope by an experimenter blind to the behavioral results.

### Data and Statistical analyses

Data are presented as means ± SEM and individual data points or box plots [medians ± 25% (percentiles) and Min/Max as whiskers], and were analyzed with STATISTICA-10 Software (Statsoft, Inc., Tulsa, OK, USA) or Statistical Package for Social Sciences (IBM SPSS, v 26, USA). Assumption for parametric analyses, namely homogeneity of variance, sphericity and normality of distribution were verified prior to each analysis with Cochran, Mauchly and Shapiro-Wilk’s tests, respectively. Where normality was substantially violated, data were Log transformed and where sphericity was violated, Greenhouse-Geisser correction was applied.

Behavioral data on acquisition or maintenance of SIPw or SIPa were analyzed using repeated measures analyses of variance (ANOVA) with either time as within-subject factor alone or time or fluid as within-subject factor and group (HD vs LD or cluster) as between-subject factor.

The effect of aDLS DA receptor blockade on adjunctive drinking was analyzed with one-way ANOVA (**Test 1**) or repeated measures ANOVA (**Test 2, 3 and 4**) with treatment as between (**Test 1**) or within-subject factor (**Tests 2-4**) and group as between-subject factor (**Tests 2-4**).

Two-step K-mean cluster analysis (58, 67) was performed to identify groups of individuals whose reliance of adjunctive drinking on aDLS DA differed across tests. Thus, fluid intake under α-flupentixol treatment at Test 2 and 4 was averaged across doses and was expressed as percentage change from baseline (i.e., vehicle-treated rats); three subpopulations of rats were identified: Cluster 1 represented aDLS reliant water coper (WC) rats (n=11), Cluster 2 encompassed the marginally aDLS reliant WC rats (n=19) and Cluster 3 the aDLS reliant alcohol coper (AC) rats (n=8).

We additionally computed an index of differential reliance on aDLS DA of compulsive drinking of alcohol vs water, using the following equation: |(sensitivity to aDLS at late SIPa - sensitivity to aDLS at late SIPw)/sensitivity to aDLS at late SIPw|. This enabled the analysis of whether the emerging reliance on aDLS control predicts individual tendency to rely on alcohol to cope with distress and the ensuing development of compulsive alcohol drinking (54).

Rats were also ranked according to their fluid intake (mL) before receiving α-flupentixol infusions (i.e., after the last three session of SIPw or SIPa) to identify individuals that increased their intake when alcohol was introduced in the SIP procedure.

The confirmation of significant main effects and differences among individual means were further analyzed using the Newman-Keuls post-hoc test, Dunnett’s test (when comparing multiple time points to a single baseline) or planned comparisons, as appropriate. Significance was set at α ≤ .05 and effect sizes are reported as partial eta squared (η_p_^2^).

## Results

Sixty rats with cannula placements located in the aDLS were included in the final analysis (**Figure 2A**).

**Figure 2:**
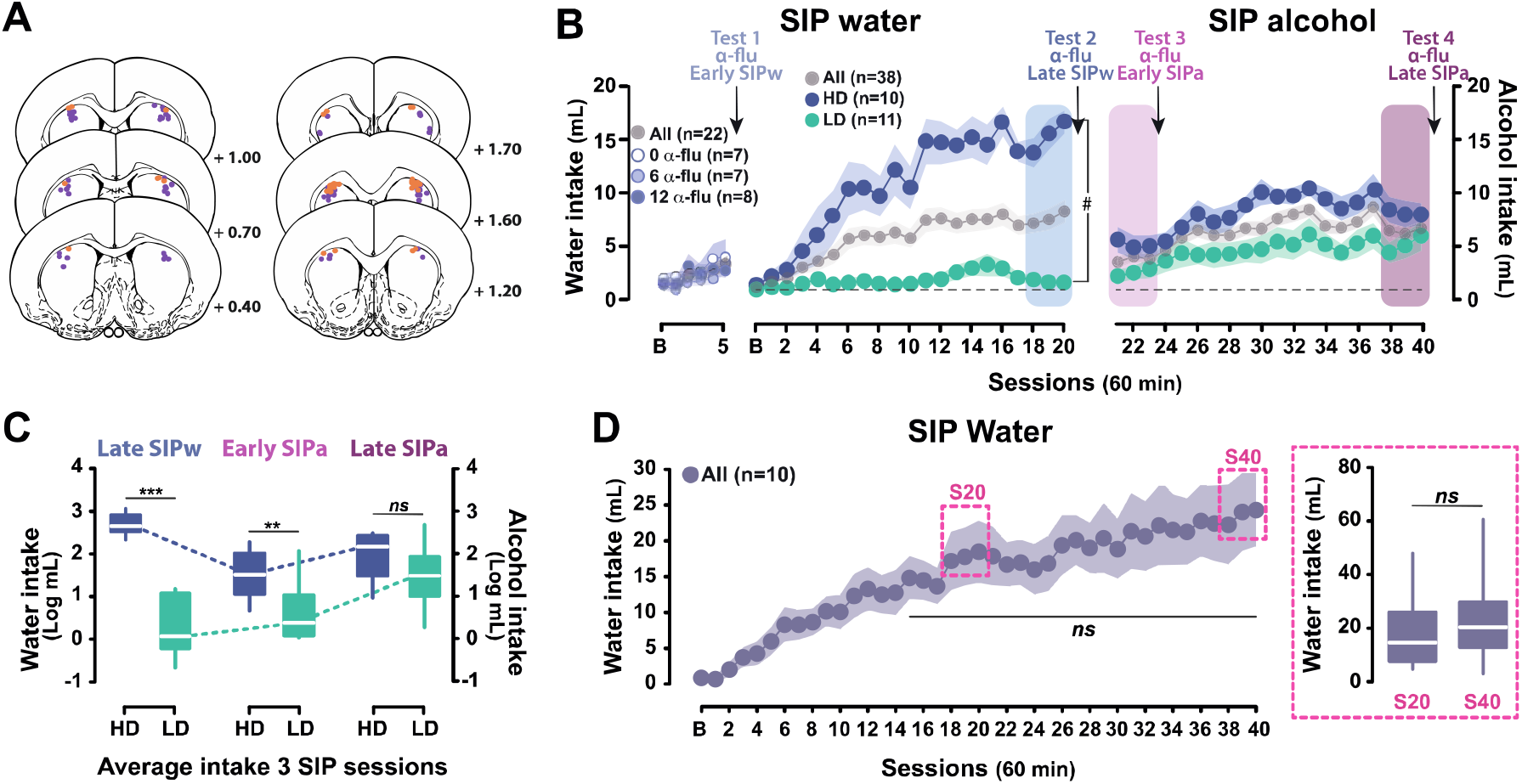
Individual differences in the tendency to develop compulsive adjunctive behaviors in rat. **A)** Rats having completed experiments 1 and 2 with cannula tips (orange and violet for experiment 1 and 2, respectively) located in the anterior dorsolateral striatum (aDLS) according to the rat brain atlas (73) as assessed following staining with Cresyl Violet were included in the final analyses. **B-C**) At the population level, and similarly across each independent group for experiment 1, food restricted rats exposed to intermittent food delivery progressively developed adjunctive polydipsic water drinking over five (Experiment 1, n=22) and twenty (Experiment 2, n=38) sessions. Experiment was designed to reveal marked individual differences in the tendency to engage in polydipsic water drinking that emerged early on in training. While at the population level individuals developed a polydipsic water drinking response, some individuals, developed an excessive, compulsive polydipsic behavior while others did not engage in a displacement strategy whatsoever. Thus, high drinker (HD) and low drinker rats (LD), selected respectively in the upper and lower quartiles of the population stratified on the average water consumption over the last 3 sessions of SIP with water (Late SIPw, blue rectangle), greatly differed in their trajectory of polydipsic water drinking. HD rats developed a compulsive water drinking behavior that reached more than 16.65 ± 1.08 ml/hour by the last session, or two times more than the whole population, whereas LD rats maintained throughout a water drinking behavior similar to that associated with their homeostatic need displayed at baseline (B). The introduction of alcohol resulted in significant changes in coping behavior. Thus, HD and LD rats still differed in their level of adjunctive drinking at the beginning of the SIP training with alcohol (Early SIPa, light purple rectangle), mostly due to the fact that HD rats, who had developed an excessive polydipsic drinking behavior with water persisted following the introduction of alcohol, albeit to a lower level. However, while HD rats maintained overall a steady level of polydipsic alcohol drinking over time, LD rats acquired a coping response with alcohol and eventually developed compulsive polydipsic alcohol drinking so that they no longer differed from HD rats by the end of training (Late SIPa, dark purple rectangle). **D**)This increase in alcohol drinking shown by LD rats was not attributable to an increase in fluid intake over time since once adjunctive drinking was established in an independent cohort of rats over 20 sessions (Experiment 3, n=10), polydipsic water drinking remained stable so that by session 40 (**S40** dashed-line rectangle, average of sessions 38-40) rats displayed a similar level of fluid intake as they did by the end of the first 20 session period (**S20** dashed-line rectangle, average of sessions 18-20). The reliance of early (Tests 1 and 3) and well-established (Tests 2 and 4) polydipsic water or alcohol drinking on aDLS DA was assessed at each time point identified by an arrow. # p<0.001 group x time interaction, *** p<0.001, ** p<0.01, HD different from LD rats, ns: no significant.

Exposure to the SIP procedure resulted, at the population level, in a progressive development of adjunctive drinking behavior, expressed as an increase in water intake in response to the introduction of a FT-60 s schedule of food delivery over 5 (for **experiment 1**) or 20 sessions (for **experiment 2**) [main effect of time: F_2.7,55.8_=5.46, p=0.003, η_p_^2^= 0.21 and F_4.4,162.4_=17.89, p < 0.001, η_p_2= 0.33, respectively] that became different from that expressed at baseline from session 4 and 3 onwards, respectively (**Figure 2B**).

As previously described (32, 54, 66, 74), marked individual differences in the propensity to lose control over adjunctive water drinking behavior as a coping strategy emerged over 20 SIPw sessions [main effect of group: F_1,19_ = 52.73, p < 0.001, η_p_^2^ = 0.73; time: F20,380 = 16.39, p < 0.001, η _p_^2^ = 0.46 and group x time interaction: F 20,380 = 11.74, p < 0.001, η _p_^2^ = 0.38] (**Figure 2B**). Thus, HD rats (upper quartile of the population, n=10) were prone readily to develop excessive levels of adjunctive water intake, eventually drinking more than 15 mL/hour (15.30 ± 1.21 mL over the last three sessions), more twice the intake of the population as a whole, and almost ten times more than LD rats (lower quartile of the population, n=11) (1.70 ± 0.32 mL over the last three sessions) [main effect of group: F_1,19_ = 113.19, p < 0.001, η _p_^2^= 0.86] (**Figure 2C**, left panel). This differential level of polydipsic water intake observed between HD and LD rats after 20 daily SIP sessions remained stable over long periods of time, as previously shown (32).

However, the introduction of the opportunity to drink alcohol instead of water as a means to cope with the distress induced by the SIP procedure resulted in an asymmetrical change in the level in adjunctive drinking displayed by LD and HD rats [main effect of time: F_2,38_ = 9.86, p < 0.001, η_p_^2^ = 0.34; group: F_1,19_ = 39.81, p < 0.001, η_p_^2^ = 0.68 and time x group interaction: F_2,38_ = 22.95, p < 0.001, η_p_^2^ = 0.55] (**Figure 2C**). Thus, despite an overall decrease in total fluid intake upon the introduction of alcohol, likely reflective of its anxiolytic properties (75), HD and LD rats initially maintained their then long established difference in the level of adjunctive drinking [HD (5.14 ± 0.84 mL/hour) vs LD (2.54 ± 0.69 mL/hour, p = 0.006] (**Figure 2C**, middle panel). However, over the 20 sessions of SIPa some LD rats (63.6%) more than doubled their intake of alcohol, eventually reaching 5.14 ± 1.12 mL per hour [LD, Early SIPa vs Late SIPa, p < 0.001] (**Figure 2C**), thereby contributing greatly to the overall increase in alcohol intake shown at the population level [main effect of time F_9.2,340_ = 12.49, p < 0.001, η^2^ = 0.25] (**Figure 2B**) while becoming as compulsive as HD rats [Late SIPa, HD vs LD, p= 0.09] that only increased their alcohol intake by ∼50% over the same period [HD, Early SIPa vs Late SIPa, p < 0.02] (**Figure 2C**, right panel). The sudden increase in fluid consumption shown by LD rats upon the introduction of alcohol cannot be accounted for by a simple increase in fluid intake over time since once established, polydipsic water intake remained stable over protracted periods of time (e.g., at least 40 days) [main effect of session: F_40,360_ = 9.49, p < 0.001, η_p_^2^ = 0.51; from session 15, between sessions all *p*-values > 0.05] (**Figure 2D**, left panel). Thus, rats exposed to the SIP procedure displayed similar level of polydipsic water intake over sessions 38-40 as over sessions 18-20 [F_1,9_ = 2.80, p = 0.13] (**Figure 2D**, right panel).

These results together replicate our original demonstration of individual differences in the reliance on alcohol to cope with distress, and the ensuing excessive drinking, which has been shown to predict and increased tendency to develop compulsive alcohol drinking (54). We then sought to identify the neural locus of control of such coping behavior that appears to be initially goal-directed, the goal of the response (drinking) being the alleviation of distress (14, 22, 76), but eventually becomes excessive and compulsive (74, 77, 78), a transition hypothesized here to reflect the development of maladaptive negative-reinforcement driven habits. Thus, we assessed the reliance of adjunctive fluid drinking on aDLS DA, a signature of habitual control over alcohol-related responding (79), and the compulsion to seek and drink alcohol (57, 58).

As predicted, after a short-term exposure to SIPw, prior to the development of excessive drinking behavior in vulnerable individuals, aDLS DA receptor blockade with α-flupentixol (0, 6, 12μg/0.5μl/side; between-subject design) had no effect on water intake [main effect of treatment: F_2,19_ <1] (**Test 1, Early SIPw, Figure 3A**). However, after extended exposure to SIPw, at a time when vulnerable individuals had developed compulsive adjunctive drinking, the same aDLS DA receptor blockade (0, 6, 12μg/0.5μl/side α-flupentixol; within-subject counter-balanced design) now resulted in a marked decrease in adjunctive water intake in these HD rats [**HD vs LD**: main effect of treatment: F_2,38_ = 4.16, p = 0.023, η_p_^2^ = 0.18; group: F_1,19_ = 44.85, p < 0.001, η_p_^2^ = 0.70 and treatment x group interaction: F_2,38_ = 2.96, p = 0.06, η_p_^2^ = 0.13]. Follow-up analyses confirmed that aDLS DA receptor blockade did not influence drinking in LD rats [main effect of treatment: F_2,20_ <1], while it greatly reduced excessive adjunctive drinking in HD rats [main effect of treatment: F_2,18_ = 3.55, p = 0.049, η_p_^2^ = 0.28; 0 vs 6 and 12µg/side: p = 0.007] (**Test 2, Late SIPw, Figure 3B**).

**Figure 3:**
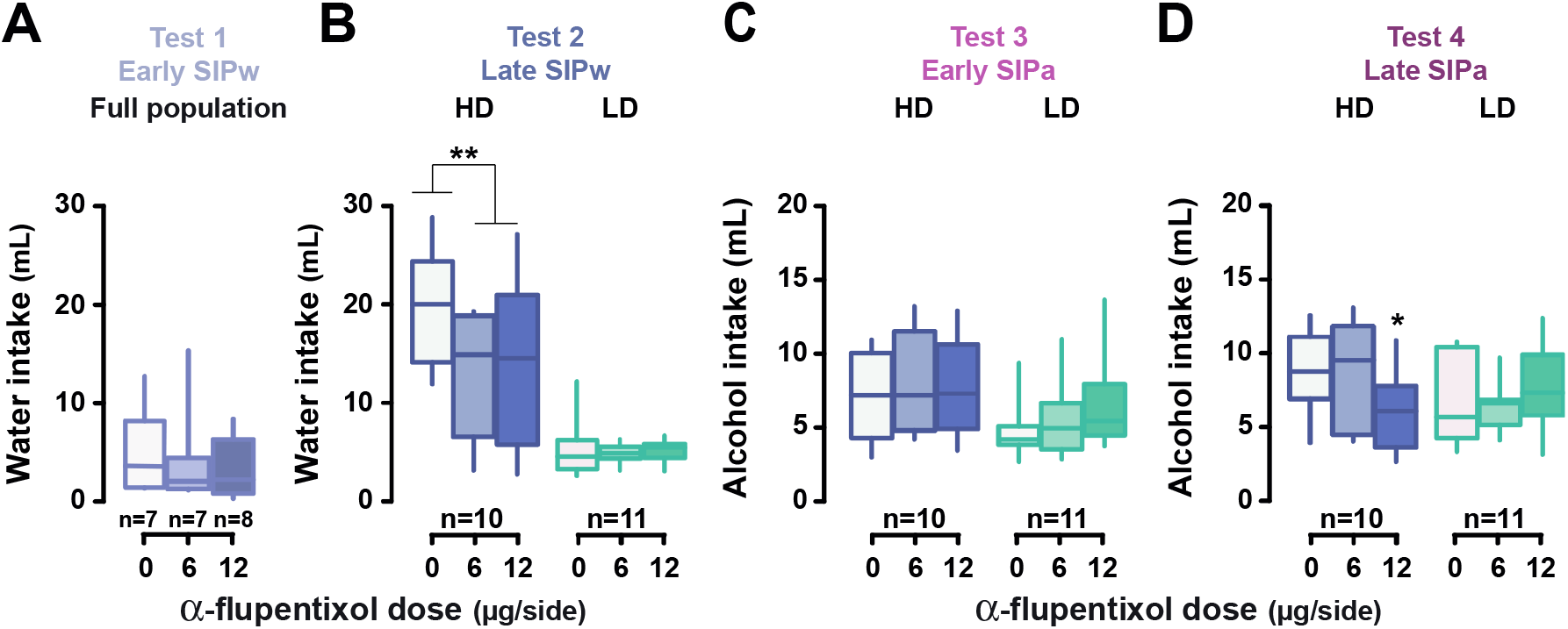
Excessive, but not early, adjunctive drinking is dependent on anterior dorsolateral striatum dopamine-dependent mechanisms. Intra-anterior dorsolateral striatum (aDLS) infusion of the dopamine receptor antagonist α-flupentixol (0, 6, 12 μg/side) had no effect on recently acquired polydipsic water intake (Test 1, **A**) or on drinking shown by low drinker (LD) rats even after 20 sessions (Test 2, **B**). In contrast, similar intra aDLS infusions of α-flupentixol (0, 6, 12 μg/side) decreased excessive water intake in high drinkers (HD) rats (Test 2, **B**). When water was replaced by alcohol, intra aDLS dopamine receptor blockage no longer influenced polydipsic drinking in HD rats while it remained ineffective in LD rats (Test 3, **C**). In marked contrast, the sensitivity shown by HD rats to intra aDLS dopamine receptor blockade when engaged in excessive polydipsic water intake re-emerged when their polydipsic alcohol intake became well established (Test 4, **D**). ** p<0.01, * p<0.05 different from vehicle (0 μg/side). SIPw: SIP with water; SIPa: SIP with alcohol.

In marked contrast, in the same rats, the introduction of alcohol as a means to cope with distress resulted in a disengagement, albeit transient (see below), of aDLS DA control over adjunctive drinking. Thus, intra aDLS infusions of the DA receptor antagonist α-flupentixol no longer decreased fluid intake in HD rats after three days of SIPa [**analyzing HD and LD**: main effect of group: F_1,19_ = 4.95, p = 0.038, η_p_2 = 0.21, treatment: F_2, 38_= 1.44, p = 0.25 and treatment x group interaction: F_2,38_ <1] (**Test 3, Early SIPa, Figure 3C**), thereby demonstrating that learning to utilize alcohol to cope with distress, even in individuals that had established a compulsive adjunctive drinking behavior with water, is associated with a disengagement of habitual control over behavior.

However, following 20 daily sessions of SIPa when adjunctive alcohol drinking had escalated and become excessive, it became, once again, reliant on aDLS DA as shown by the emergence of a sensitivity of alcohol drinking to aDLS DA receptor blockade in HD rats [**analyzing HD and LD**: main effect of group: F_1,19_ = 1.17, p = 0.29 and treatment x group interaction: F_2,38_ = 4.76, p = 0.014, η_p_^2^ = 0.20; HD rats, 0 vs 12 μg/side: p = 0.044] (**Test 4, Late SIPa, Figure 3D**).

Building on previous evidence that the magnitude of the reliance of instrumental alcohol seeking on aDLS DA when it becomes habitual predicts the severity of the ensuing compulsive behavior in vulnerable individuals in a positive reinforcement setting (58), here we further investigated whether a similar relationship was observed between the magnitude of the reliance of adjunctive responses, maintained through negative reinforcement, on aDLS DA and the vulnerability to develop excessive, compulsive adjunctive drinking.

We capitalized on the within-subject approach on which experiment 2 was designed to determine, for each individual, the differential sensitivity of excessive adjunctive drinking of water (**Test 2**) vs alcohol (**Test 4**) to intra aDLS infusions of α-flupentixol (mean % decrease of intake following DA receptor blockade relative to baseline, i.e., following vehicle infusions). A K-mean cluster analysis (57, 67) revealed three sub-populations that differed in the reliance of their excessive water or alcohol adjunctive drinking on aDLS DA (**Figure 4A**). A first cluster (Cluster 1, 28.9% of the total population) comprised individuals that had all developed an aDLS-DA dependent adjunctive water drinking behavior before they switched to alcohol drinking that also became reliant on aDLS-DA. These aDLS reliant WC rats, which comprised 82% of HD and intermediate individuals, differed from rats of Cluster 2 (50% of the population; which instead comprised 84% of intermediate and LD rats) whose water drinking was marginally reliant on aDLS-DA and who did not engage their habit system when subsequently exposed to alcohol. In contrast, individuals in Cluster 3 (21.1% of the population, which comprised 87.5% of LD and intermediate rats) were the only ones to develop an adjunctive response reliant on aDLS DA when they used alcohol, as confirmed by an analysis of the ratio of sensitivity of adjunctive fluid intake to aDLS DA receptor blockade between SIPa and SIPw [main effect of cluster: F_2,35_ = 13.33, p < 0.001, η_p_^2^ = 0.43; Cluster 3 vs 1 and 2, p < 0.001, respectively) (**Figure 4B**, bottom right panel). The time course of the engagement of aDLS DA in the control over adjunctive behavior of Cluster 3 rats was therefore very different to that shown by Cluster 1 rats (**Figure 4C**) [treatment x time x cluster interaction: F_5,87.3_ = 4.48, p = 0.001, η_p_^2^ = 0.20; treatment x time interaction: F_2.5,87.3_ = 4.48, p = 0.005, η_p_^2^ = 0.12], with profound differences in their sensitivity to aDLS DA receptor antagonism at LATE SIPw (**Test 2**) [main effect of treatment: F_1.5,52.2_ = 5.01, p = 0.017, η_p_^2^ = 0.12; treatment x cluster interaction: F3,52.2= 6.16, p = 0.001, η_p_^2^ = 0.26] and at LATE SIPa (**Test 4**) [main effect of treatment: F_1.7,60.1_ = 6.49, p = 0.004, η_p_^2^ = 0.16; treatment x cluster interaction: F_3.4,60.1_ = 7.55, p < 0.001, η_p_^2^ = 0.30], but not EARLY SIPa (**Test 3**) [main effect of treatment: F_2,70_ = 1.22, p = 0.30; treatment x cluster interaction: F_4,70_ <1]. The differences at LATE SIPw were driven exclusively by Cluster 1 individuals [0 vs 6 μg/side: p < 0.001, 0 vs 12 μg/side: p < 0.001] (**Figure 4C**, upper panel), whereas the effects at LATE SIPa were driven by individuals belonging to both Cluster 1 and Cluster 3 [**Cluster 1**, 0 vs 12 μg/side: p = 0.001; **Cluster 3**, 0 vs 12 μg/side: p = 0.010] (**Figure 4C**, bottom panel). Further retrospective investigation of the individuals comprised in each cluster revealed that most of those belonging to Cluster 1 were HD rats that had developed a compulsive adjunctive drinking behavior with water which then persisted when alcohol was introduced, whereas the majority of individuals in Cluster 3 instead had developed compulsive adjunctive fluid intake only upon introduction of alcohol. These individuals were actually AC that increased their polydipsic alcohol intake by 60% over the three weeks of SIPa, eventually to reach the level of intake shown early on by WC soon after the introduction of alcohol. Together these data demonstrate that the development of compulsive adjunctive drinking depends on the engagement of aDLS-DA dependent control over behavior that eventually occurred in 81.6 % of the individuals in this study.

**Figure 4:**
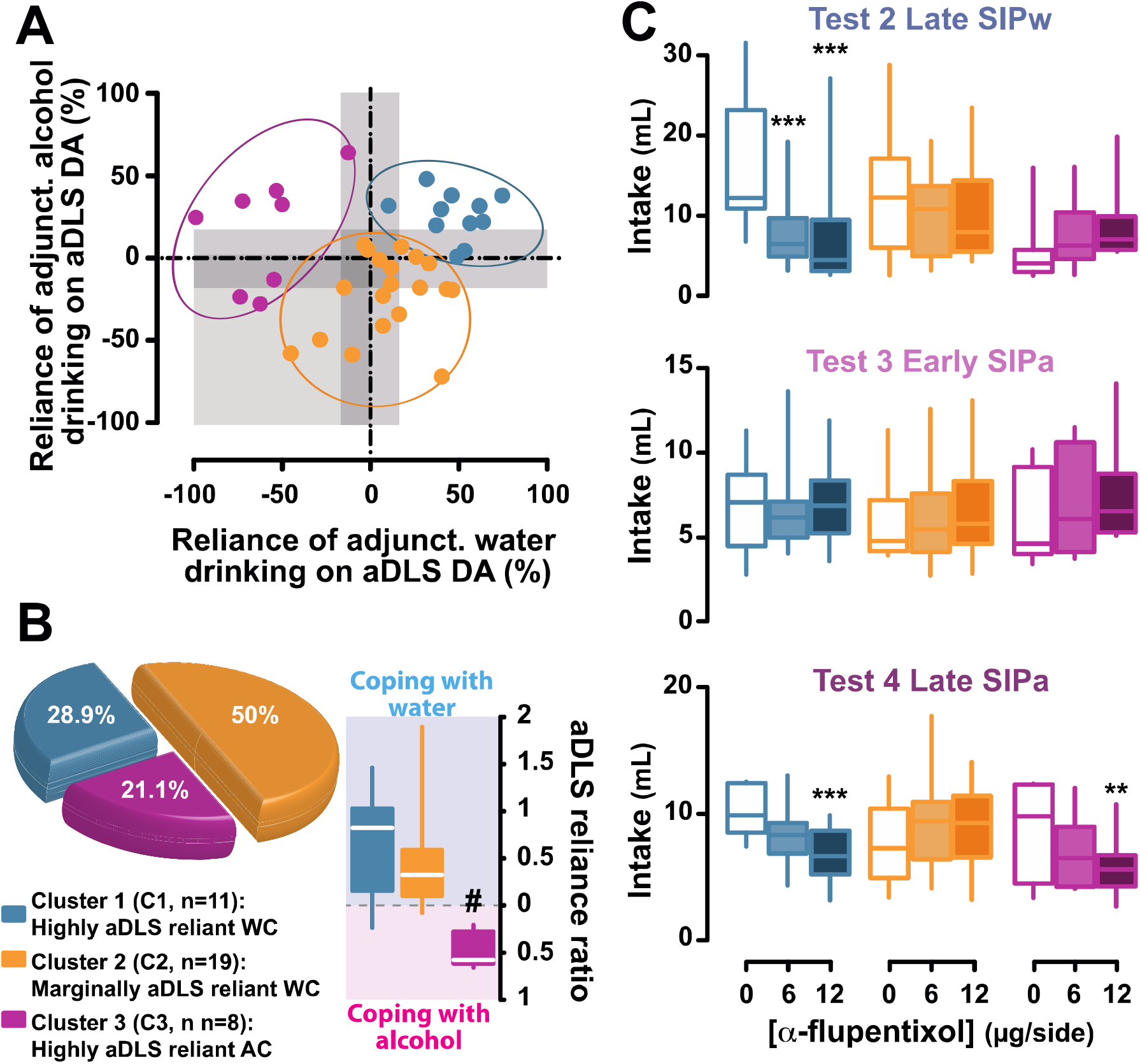
The individual tendency to develop anterior dorsolateral striatum dopamine-dependent-coping response is associated with the vulnerability to develop compulsive adjunctive behavior. **A-B**) Marked individual differences in the reliance of compulsive drinking on anterior dorsolateral striatum (aDLS) dopamine (DA) were revealed when normalizing the influence of aDLS DA receptor blockade on polydipsic water (Late SIP water) or alcohol intake (Late SIP alcohol) to the baseline levels of drinking following vehicle infusions (0 μg/side). A cluster analysis identified three subpopulations of rats, one (Cluster 1) comprised individuals whose polydipsic behavior, irrespective of the fluid drank, was heavily reliant on aDLS DA. These highly aDLS reliant water copers (WC) represented 28.9% of the overall population and consisted predominantly (82%) of high drinkers (HD) and intermediate individuals. A second cluster (Cluster 2) comprised individuals whose polydipsic water or alcohol drinking behavior was overall marginally sensitive to aDLS DA receptor blockade. These marginally aDLS reliant WC represented 50% of the population and consisted predominantly (84%) of low drinkers (LD) and intermediate rats. The third cluster (Cluster 3) comprised individuals whose polydipsic alcohol drinking was much more reliant on aDLS DA than their polydipsic water drinking behavior. These highly aDLS reliant alcohol coper rats (AC) represented 21% of the population and consisted predominantly (87.5%) of LD and intermediate rats. A systematic analysis of the respective reliance of polydipsic water vs alcohol drinking behavior on aDLS DA confirmed that only highly aDLS reliant AC displayed a selective increased in the engagement of aDLS DA-dependent control over behavior when they used alcohol as a mean to cope with distress. **C)** Marked differences were observed in the time course of the engagement of the aDLS-DA dependent habit system in mediating compulsive coping behavior in the individuals of these three groups that map perfectly those observed on HD and LD rats (Figure 3), in that rats from cluster 1 showed sensitivity to aDLS DA receptor blockade when SIPw (Test 2 Late SIPw) and SIPa (Test 4 Late SIPa) were well-established whereas rats from cluster 3 showed such sensitivity only when SIPa was well-established, neither showing reliance on aDLS DA during the acquisition of SIPa. Rats from the heterogenous cluster 2 never showed a decrease in adjunctive drinking following aDLS that reached statistical significance throughout the training history. *** p≤0.001, ** p≤0.01, compared with the vehicle treatment (0 μg/side); # p<0.001 compared with cluster 1 and 2.

## Discussion

At the population level, individuals in the present study developed an adaptive coping response to the distress generated by intermittent delivery of food under a SIP procedure, expressed as non-regulatory, adjunctive polydipsic drinking behavior that remained stable over up to 42 days. However, within two weeks some individuals progressively lost control over the polydipsic drinking response and developed a compulsive coping strategy. In contrast, some individuals did not acquire a coping response with water, but only did so when they had access to alcohol. These individuals that drank alcohol to cope with distress subsequently developed compulsive coping behavior.

The results of the present study replicate previously reported individual variability in the tendency both to engage in polydipsic drinking (54) and to develop compulsive adjunctive behaviors (32, 54, 66, 68, 69, 74, 77, 78, 80), and more recent evidence that some individual rats that do not develop a coping response with water, do so readily with alcohol (54). Importantly, the results of the present study provide causal evidence of a progressive engagement of aDLS DA in the control over adjunctive behavior, especially when it becomes compulsive in vulnerable individuals. While the acquisition of polydipsic drinking, of water or alcohol was not dependent on aDLS DA, its excessive and compulsive manifestation in vulnerable individuals was selectively decreased by bilateral aDLS DA receptor blockade.

In those individuals that had not developed a coping strategy with water (so-called AC, 54) polydipsic drinking became reliant on aDLS DA only when alcohol drinking became compulsive. This engagement of aDLS DA habit system (81, 82) in the control over polydipsic alcohol drinking was specific to a facilitated transition to compulsion by the acquisition of alcohol use as a self-medication strategy. It could not be attributed to the instantiation of habitual control over behavior by protracted training (83) because (i) the individual differences observed in the tendency to develop excessive polydipsic drinking after 20 sessions of SIPw did not evolve further across an additional period of 20 sessions when animals continued to be given access to water instead of alcohol (see **Figure 2D** and 32), and (ii) some individuals that had not engaged aDLS DA when drinking under SIPw also did not do so when subsequently given access to alcohol (**Figure 3**). Finally, the emerging reliance on aDLS DA of alcohol drinking as a coping behavior cannot be explained by a difference in the level of fluid intake: while the level of compulsive fluid intake shown by HD rats at the end of the 20 sessions of SIPa was much lower than that shown at the end of the 20 sessions of SIPw, in both cases their excessive adjunctive behavior was equally sensitive to aDLS DA receptor blockade. Importantly, although they decreased the volume of fluid they drank when alcohol was introduced, HD rats still ingested 5-8 ml/hour, that is more than double the amount of alcohol consumed by the same strain of rats (1-3 ml/h) following at least 12 sessions of intermittent access to 10% alcohol in a two-bottle choice procedure (data from 54)), a time when rats had already escalated their alcohol intake. These observations reveal the excessive nature of their alcohol intake, which has previously been shown to result in blood alcohol levels higher than 0.6 g.l^-1^ (54). These results are in line with a body of evidence suggesting marked differences in the psychological and neural mechanisms underlying the acquisition of adjunctive coping responses and its subsequent compulsive manifestation in vulnerable individuals.

The psychological nature of adjunctive behaviors has long been debated since they were originally considered not to fulfil the necessary requirements of contingency with food delivery to be instrumental in that polydipsic drinking, even in the early acquisition stages, is a consequence of the intermittent delivery of food. It does not lead to the delivery of food or help to meet an homeostatic need (for review, see 76).

However, if a change in an interoceptive state, namely anxiolysis (84), is considered the outcome of the coping displacement strategies, then adjunctive polydipsic drinking, which superficially might appear to be consummatory, is in fact an instrumental response (76, 85, 86), elicited and maintained by negative reinforcement. Furthermore, the engagement in polydipsic drinking during the first week of exposure to SIP depends on the interoceptive cortex, namely the anterior insula (32) and results in a decrease in the activation of the hypothalamic-pituitary-adrenal axis provoked by intermittent food delivery (14, 15, 19-22). Furthermore, at the same early stage of training, prevention of drinking in the SIP context results in an increase in corticosterone levels (13).

Early polydipsic water intake has features of goal-directedness in that it is sensitive to outcome devaluation (anti-anxiolysis), since there is a decrease in responding following administration of anxiogenic drugs such as CRF (87) or amphetamine (63, 88). In contrast, following prolonged exposure to SIP, the same acute amphetamine challenge no longer results in a decrease in drinking (88). These observations suggest that polydipsic drinking is initially a response tied to the motivational value of its outcome, namely anxiolysis, but that it eventually becomes habitual (89).

Development of habitual control over polydipsic drinking is consistent with a progressive shift in the striatal locus of control over behavior. Selective 6-hydroxydopamine lesions of the mesolimbic system prevent the acquisition of SIP (90-92), which is otherwise associated with a ramping of extracellular levels of DA in the nucleus accumbens core across sessions (93). In contrast, DA transporter deficient mice and rats, which have elevated levels of DA, display an impaired acquisition of polydipsic drinking (94, 95), which is also decreased in wild type animals by the acute administration of psychostimulants such as cocaine and D-amphetamine directly into the nucleus accumbens (31, 96). Together these results show that the acquisition of adjunctive drinking behavior depends on DA signalling in the mesolimbic system, but that it is impaired by an increase in tonic extracellular levels of DA. In marked contrast, when this coping behavior is well established and becomes excessive in vulnerable individuals, it is no longer impaired by similar causal manipulations of the mesolimbic DA system (63). Furthermore, repeated amphetamine exposure, which facilitates habitual responding under positive reinforcement (97), exacerbates polydipsic drinking (98) which becomes independent of food delivery (98, 99), suggesting the development of stimulus-response control over behavior (100) when it has become compulsive in vulnerable individuals. These observations are in line with the evidence that polydipsic drinking during early SIPw training is not accompanied by neuronal activation in the DLS (101), whereas its compulsive manifestation after extended exposure is associated with increased spine density in this region (64).

The apparent ventral to dorsal striatal shift in the locus of control over adjunctive behaviors when they become compulsive (102) agrees with previous evidence that the development of compulsive alcohol seeking in vulnerable individuals is predicated on the functional engagement of aDLS DA. This lends further support to the hypothesis that the development of compulsion stems from a loss of control over aDLS dependent habits (103-106). It has been recently shown in the context of cocaine addiction (70) that compulsive coping behavior may be conceptualized as the manifestation of an urge to express an ingrained habit (105, 107, 108) the performance of which actually becomes the goal of the behavior (70).

While stress shifts the balance towards aDLS-dependent habits from goal-directed behavior (109-111), it is unlikely that the individual differences in the tendency to develop compulsive polydipsic water drinking observed here and elsewhere (32, 66, 68, 69) are due to a differential sensitivity to stress. Individuals that did not acquire a coping response with water did so readily when they had access to alcohol instead. This suggests that the lack of development of an adjunctive behavior under SIPw in these individuals was not due to their lack of sensitivity to the distress engendered by the SIP procedure. Instead, these individuals needed alcohol in order to develop this coping response.

In contrast, the tendency of WC to develop a coping response with water and the subsequent loss of control over it in vulnerable individuals may reflect an interplay between a heightened sensitivity to negative urgency (112) and an inherent propensity generally to rely on the aDLS habit system. Hence, on the one hand, the development of compulsive SIP depends on both insula-dependent interoceptive mechanisms, the noradrenaline stress system and their interaction with pre-existing impulsivity (32, 66). Whereas on the other hand, it is also predicted by a pre-existing tendency to use the habit system across a wide array of tasks, from response learning in spatial navigation (113, 114) to resistance to devaluation in instrumental reinforcement and perseverative responses in reversal learning task (101, 115). Overall, these results support the view that the inflexibility and increased responding under operant schedules that promote habitual behavior shown by HD for water rats (69, 116) arises from an interaction between an inherent tendency to rely on the habit system and its recruitment by negative reinforcement during the development of a coping response.

Using the same SIP procedure, we have previously shown that alcohol enables the development of adjunctive responses in a specific subpopulation of rats otherwise unable to cope with distress by drinking water. In these individuals, it was the acquisition of alcohol use as a self-medication strategy, and not the overall level of intoxication (AC did not differ from WC in their overall level of alcohol intake or the blood alcohol levels they reach) that determined the greater vulnerability they showed to develop compulsive, quinine resistant, alcohol drinking (54). This subpopulation of AC is very similar to that identified as Cluster 3 in the present study which only engaged aDLS DA when alcohol was introduced in the SIP procedure. The engagement of aDLS DA in these AC may be attributable to the adaptations that alcohol exposure causes in the aDLS including its disinhibition by the dampening of GABAergic transmission onto its principal medium spiny projection neurons (117, 118). Together, these observations support the view that the engagement of habitual control over negative-reinforcement driven behaviors, such as the acquisition of alcohol drinking as a self-medication strategy, is an important determinant of the vulnerability to develop compulsive drinking (52, 56, 104, 119-121).

It remains to be established why some individuals can only develop a coping response with alcohol but not water. One hypothesis is that the anxiolytic properties of alcohol facilitate the acquisition of the adjunctive response in these individuals (122, 123). This is suggested by the marked and steady decrease in total fluid intake WC show after introduction of alcohol, which is quantitatively similar to that observed following administration of benzodiazepines (124).

Importantly, the introduction of alcohol in these WC resulted in a transient disengagement of the aDLS DA-dependent habit system. This reveals that the same behavioral expression of a coping response, similar to instrumental responses for positive reinforcers, can be mediated by either the goal-directed or the habit system. But perhaps more importantly it suggests that even a well-established, negative reinforcement-based habitual coping response can undergo incentive learning (89, 125) when alcohol becomes part of it.

Together these observations provide evidence for a role of negative reinforcement-based habits in the development of compulsive coping behaviors.

## Acknowledgements

This work, carried out at the department of Psychology of the University of Cambridge, was funded by a UKRI grant (MR/N02530X/1) to BJE and DB, Brain and Behavior Research Foundation NARSAD Young Investigator Grant (RG95599 Grant ID: 27353) to CG, and Leverhulme Trust Early Career (ECF-2018-713) and Isaac Newton Trust fellowship (18.08(g)) to LMP.

For the purpose of open access, the author has applied a Creative Commons Attribution (CC BY) licence to any Author Accepted Manuscript version arising.

The data can be available at: https://drive.google.com/drive/folders/1_qrEP-ztVDG3IOMR6Ytza5cKnCbbVvzn?usp=sharing

We authors would like to thank Drs Aude Belin-Rauscent, Antonio Ferragud and Clara Velazquez-Sanchez, as well as Laetitia Ward, for technical assistance with behavioral training

## Authors contribution

DB, CG, AD, and YPO designed the experiments. CG, AD, YPO, MP and CM carried out the behavioral experiments. LMP, CG and DB performed the data analysis. LMP, CG, BJE and DB wrote the manuscript.

## Notes

**Conflict of Interest:** The authors declare no competing financial interests.

### Competing Interest Statement

The authors have declared no competing interest.

